# Information-rich localization microscopy through machine learning

**DOI:** 10.1101/373878

**Authors:** Taehwan Kim, Seonah Moon, Ke Xu

## Abstract

While current single-molecule localization microscopy (SMLM) methods often rely on the target-specific alteration of the point spread function (PSF) to encode the multidimensional contents of single fluorophores, we argue that the details of the PSF in an unmodified microscope already contain rich, multidimensional information. We introduce a data-driven approach in which artificial neural networks (ANNs) are trained to make a direct link between an experimental PSF image and its underlying parameters. To demonstrate this concept in real systems, we decipher in fixed cells both the colors and the axial positions of single molecules in regular SMLM data.

---

Originally developed toward the specific goal of superior spatial resolution, single-molecule localization (super-resolution) microscopy (SMLM), including STORM/(F)PALM^1–3^ and PAINT^4^, has evolved in recent years into a generic tool for sampling diverse biologically relevant information at the nanoscale^5^. For example, when combined with environment-sensitive dyes, intracellular heterogeneity in the local chemical environment can be mapped by concurrently obtaining the spatial positions and emission spectra of millions of single fluorescent molecules^6^. Consequently, the full potential of SMLM is expected to be unleashed through the proper multidimensional analysis of the emission wavelength^5^, brightness, dipole orientation^7^, as well as the axial (3D) position^8^ of single molecules.

To date, to extract information beyond the in-plane location, e.g., the emission wavelength and the axial position, of single emitters, would often oblige the explicit encoding of such high-dimensionality information into the diffraction pattern of single molecules (point spread functions; PSFs) through optical aberrations and alterations^5,8^, including astigmatism^9^, interference^10^, wavelength-dependent splitting^11^, dispersion^12^, and wave-front modification^13^. The resultant, engineered PSF shape and intensity then help establish best-fit models between experimental observables and fluorophore characteristics. Such approaches, each often optimized for a single parameter of interest, inevitably increase the PSF size and/or necessitate the splitting of fluorescence across different channels, and so often incur complicated optics and compromised performances between different parameters. While recent work^14,15^ has studied the PSF design for the simultaneous estimation of color and axial position, added optics and enlarged PSFs are still involved, and proper calibration of such Fourier optics-heavy systems is challenging^16^.

We reason that even the simplest PSF obtained from an unmodified microscope is rich in information—in addition to the axial location embedded in the defocused PSF, which has been examined in recent work^17,18^, the emission wavelength of a fluorophore also sets the scale of PSF in all three dimensions^19^. Contributions from the two sources are distinct yet subtle, and would be hard to decouple via simple models given the difficulties in fully characterizing all system-specific properties.

We present a data-driven approach in which the relationship between a PSF image, obtained from an unmodified commercial microscope, and the underlying multidimensional characteristics of an emitter is directly established by a supervised machine learning algorithm. A related approach has been recently used in astronomy for stellar classification^20^. Although SMLM faces additional challenges associated with the vast range of axial positions (as opposed to stars always at infinity), it benefits from the ready access to arbitrary amounts of experimental PSFs that may be acquired under identical conditions. By training generic learning models using such datasets, an end-to-end framework from raw PSF images to the molecule characteristics can be constructed.

To demonstrate this concept, we developed a method for machine learning-based 3D multi-color SMLM (Fig. 1 and **Methods**). With typical experimental pixel sizes (~100 nm), the dimensionality of the PSF images is moderate (modeled as 13×13 pixels), and thus artificial neural networks (ANN) with multiple hidden layers^21^ were directly used as our learning model. ANN is beneficial here as it possesses excellent representational power; moreover, since ANN does not require domain-specific expertise to construct nonlinear models, it offers a facile means to train different target parameters in an end-to-end fashion.

**Fig. 1.**
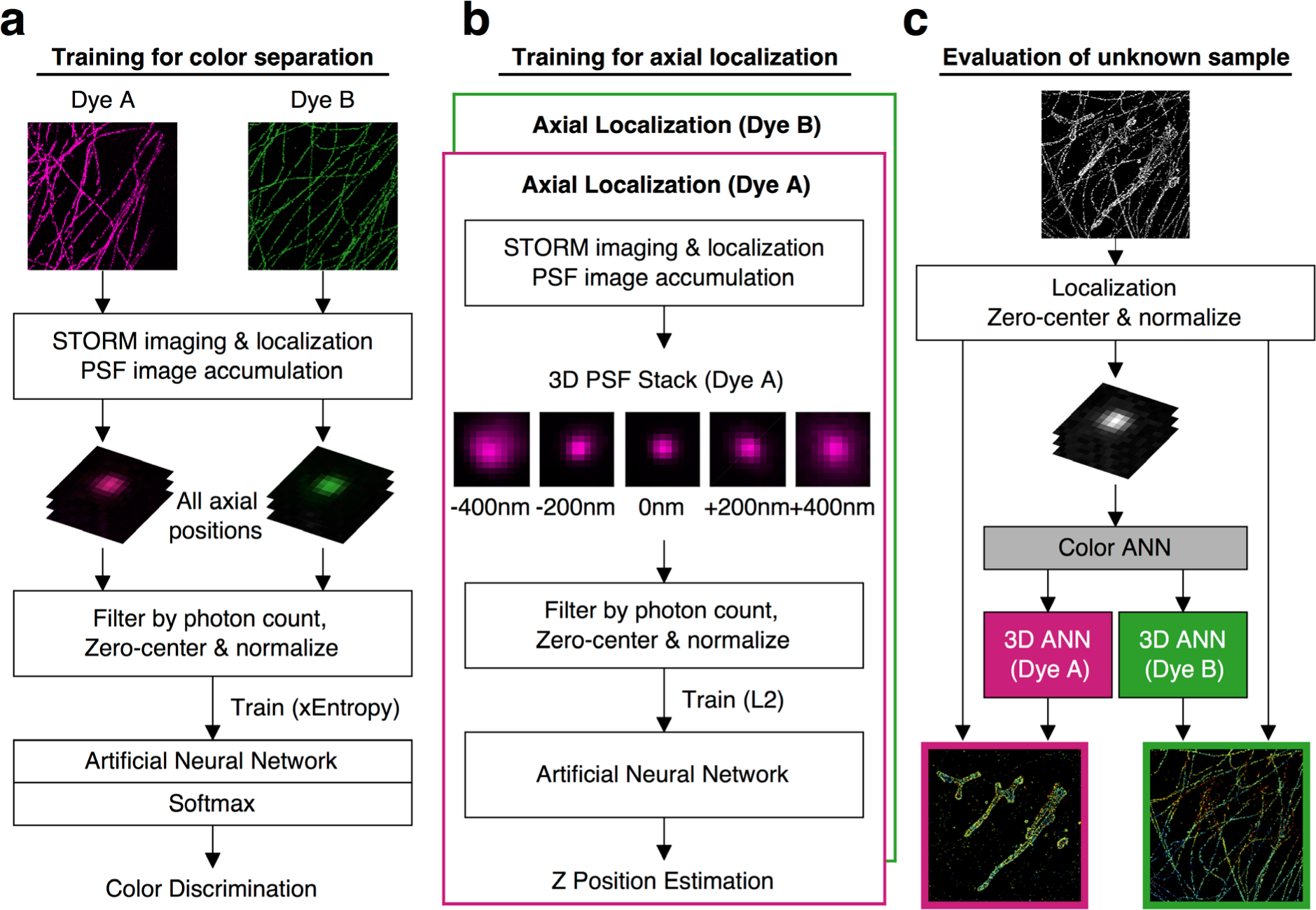
Workflow of the machine learning-based multidimensional SMLM. **(a)** A color-separating ANN is trained using samples each singly labeled by one known fluorophore, in which PSFs at different axial positions are well-represented. **(b)** ANNs for resolving the axial position are separately trained for each fluorophore using PSFs of known axial positions. **(c)** For the analysis of unknown samples, single-molecule images are localized in 2D, and first fed into the color-separating ANN described in (**a**). The color-separated single-molecule images are then separately fed into the axial-localization ANNs trained with the corresponding fluorophores, as described in (**b**). The resultant color and axial position information is then combined with the 2D localization of each molecule to generate the final multidimensional SMLM data.

One ANN with a final Softmax layer was first trained using cross-entropy loss to determine the emitter color of each PSF. Once trained, the final Softmax output provided an estimate for the conditional probability distribution of the fluorophore color, which enabled the classification of each PSF image with known confidence (Fig. 1a and **Methods**). For this color-separating ANN, training data for different fluorophores were separately prepared from multiple imaging sessions performed under the same experimental conditions as the final sample, but using only one known fluorophore at a time. The training data contained sufficient samples for fluorophores at different axial positions within the depth of field (~±500 nm of the focal plane), so that the ANN was trained to recognize fluorophores for all axial positions.

In parallel, ANNs for resolving the axial position of the emitter were separately trained for each fluorophore using L2 loss so that the final output was a scalar value^22^ corresponding to the decoded axial position (Fig. 1b). Training data for these axial-localization ANNs were collected by step-scanning samples each containing one specific fluorophore, as is typically performed for the calibration in existing 3D SMLM methods^9^.

Once both trainings were completed, SMLM data from unknown samples were localized in 2D, and the single-molecule images were first fed into the above color-separating ANN (Fig. 1c). The resultant, color-separated single-molecule images were then separately fed into the above axial-localization ANNs trained with the corresponding fluorophores (Fig. 1c). Multidimensional SMLM data were thus obtained by integrating the ANN-inferred color and axial information with the initial 2D-localization results.

We first examined the performance of the color-separating ANN using two types of 40-nm dia. fluorescent beads that differed by 45 nm in emission wavelength (“yellow” and “orange”). When the beads are at the focal plane, fitting to simple Gaussian models yielded PSF sizes (2σ) that were directly proportional to the emission wavelength (Fig. 2a), as expected. At photon counts of 3000-4000, this difference gave adequate separation of the two colors (Fig. 2a). However, this separation quickly fell apart when results from different axial positions were mixed: unsurprisingly, defocusing led to substantially increased PSF sizes, and so this parameter no longer offers useable color separation (Fig. 2b).

**Fig. 2.**
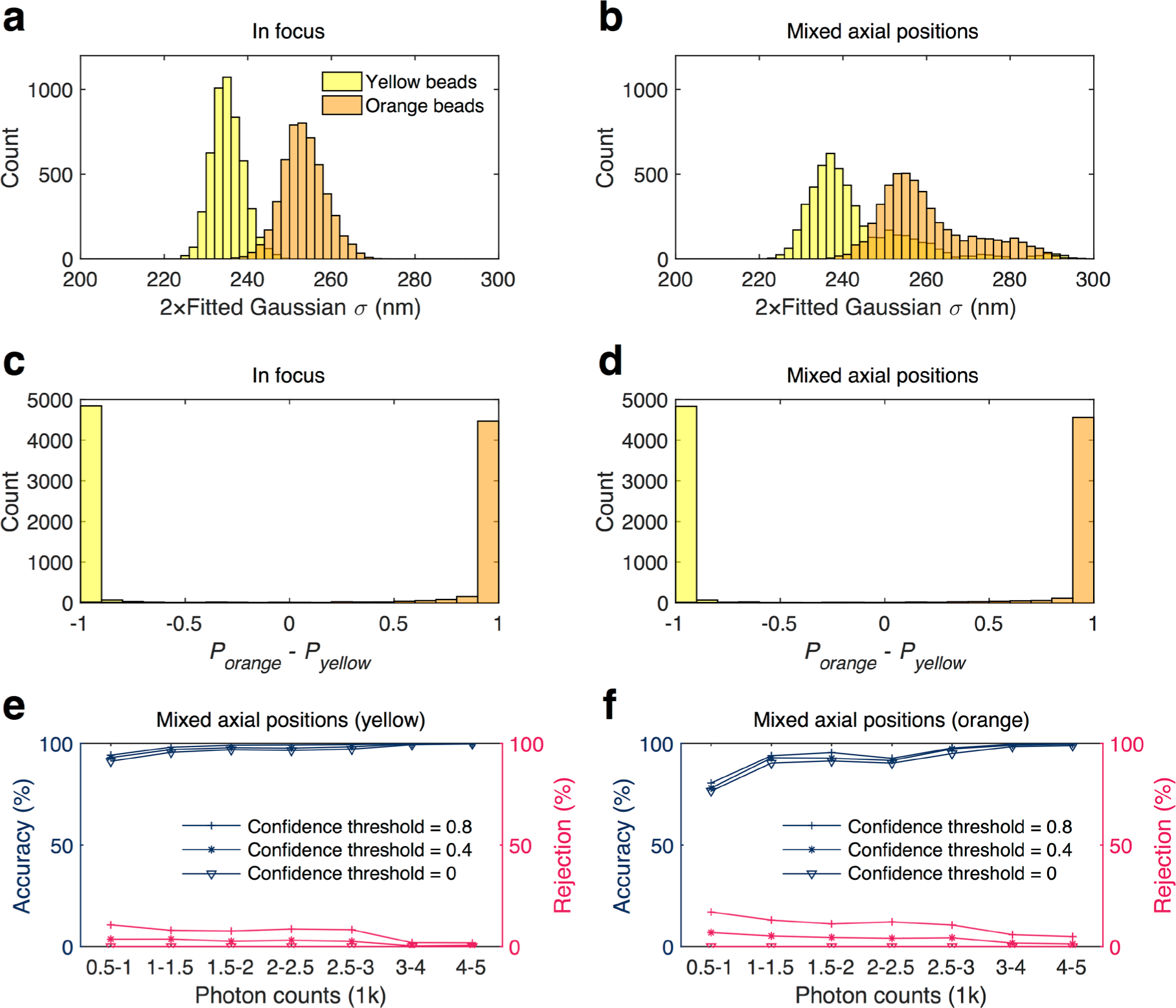
Performance of the color-separating ANN for fluorescent beads. (**a-b**) PSF size (2σ of 2D Gaussian fitting) distribution of “yellow” and “orange” beads emitting 3000-4000 photons, for (**a**) all beads are right at the focal plane and (**b**) the focus is evenly scanned over ±200 nm of the focal center. (**c**-**d**) Outputs of our color-separating ANN on the same datasets, presented as the distribution for the differences in the evaluated probabilities of each bead being “orange” vs. being “yellow”. (**e-f**) Accuracy of classification (left axes) and rejection rate (right axes) as a function of photon count for the “yellow” (**e**) and “orange” (**f**) beads in the presence of defocusing, for confidence thresholds of 0, 0.4, and 0.8.

In contrast, our color-separating ANN recognized the nuances in the PSF patterns due to differences in color vs. differences in axial position, and thus offered excellent color separation both in the presence and absence of defocusing (Fig. 2cd). As mentioned, the output of this ANN gives the conditional probabilities of each given single-molecule image being classified as certain types of fluorophores. In the binary yellow-orange system, the results can be simplified as the difference *Δ* between the evaluated probabilities of being orange and being yellow for every image (Fig. 2cd). Even in the presence of defocusing, simple classification based on *Δ*>0 and *Δ*<0 gave near-perfect (~99% accuracy) identification for both types of beads in the photon count range of 3000-4000 (Fig. 2ef). Note in STORM experiments, an average of >5000 photons is often obtainable for single molecules^9,23^. Reducing the photon count to the range of 1000-1500 photons led to modest decreases in accuracy to 94.9% and 89.1% for the yellow and orange beads, respectively (Fig. 2ef). These results were improved to 97.9% and 93.1%, respectively, by only keeping classifications with |*Δ*| above the confidence threshold of 0.8, at the expense of rejecting 8.6% and 13.5% classifications for the yellow and orange beads, respectively (Fig. 2e). Our ANN approach can thus be readily tuned for experiments that emphasize color-separation accuracy vs. experiments that emphasize the retention of molecules.

We next tackled cell imaging (Fig. 3). The microtubules and mitochondria in adherent COS-7 cells were respectively immunolabeled with two STORM dyes, CF568 and Alexa Fluor 647 (AF647). Both dyes were excited within the same STORM imaging session, and resultant single-molecule fluorescence was collected in one single optical path after a multi-notch filter. For training of the first, color-separating ANN, COS-7 cells singly-labeled by CF568 and AF647 for microtubules were STORM-imaged on the same setup, which naturally contained single molecules at all possible axial positions within the depth of field. For training of the second, axial-localization ANN, dye-labeled antibodies were attached to the coverslip for step-scanning in the axial direction (**Methods**).

**Fig. 3.**
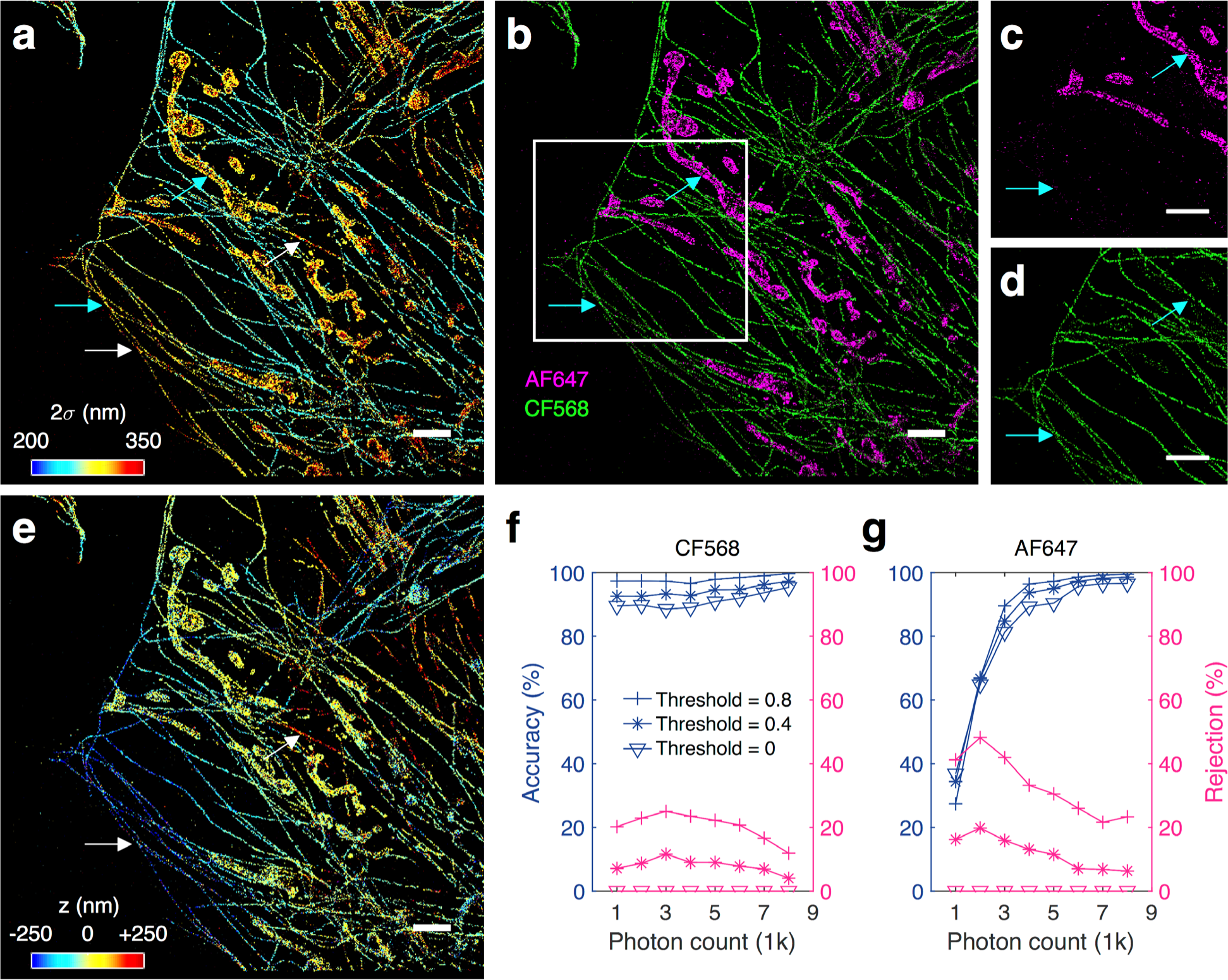
ANN-resolved multicolor 3D SMLM in cells based on unmodified PSFs. (**a**) STORM image of CF568-labeled microtubules and AF647-labeled mitochondria in a fixed COS-7 cell, colored by the fitted Gaussian width (2σ) of the PSF of each molecule, for molecules brighter than 3,000 photons. **(b)** Result of the color-separating ANN for the same dataset, at a confidence threshold of 0.8. (**c-d**) The separated AF647 (**c**) and CF568 (**d**) channels for the boxed area in (**b**). Cyan arrows in (**a-d**) point to two regions where molecules of similar PSF widths are correctly determined as different colors by the ANN. **(e)** The merged 3D STORM image after separately determining the axial position of every single molecule based on ANNs respectively trained for AF647 and CF568. Color here presents the axial position (*z*), with blue being closest to the substrate and red being the farthest away. White arrows in (**a)** and (**e**) point to two regions of the CF568 microtubule labeling that showed similar defocusing effects, but determined by ANN as being on opposite sides of the focal plane. Scale bars, 2 μm (**a-e**). **(f-g)** Classification accuracy (left axis) and rejection rate (right axis) of the color-separating ANN as a function of photon count, for cells singly labeled by CF568 (**f**) and AF647 (**g**), at confidence thresholds of 0, 0.4, and 0.8.

Figure 3a presents the acquired STORM image colored by the fitted Gaussian width (2σ) of the PSF of each molecule. Whereas it is clear that all the narrowest widths belonged to microtubules, which were stained by the shorter-wavelength dye CF568, larger widths were found at both microtubules and mitochondria (e.g., cyan arrows in Fig. 3a). This result is similar to what we saw in the bead data (Fig. 2b): defocusing broadens the PSF width, and so this simple parameter can no longer be used to separate colors. Remarkably, our color-separating ANN achieved excellent color separation for the entire image independent of axial position (and thus defocusing) (Fig. 3b-d), and consistent results were obtained on different cells over repeated experiments (Supplementary Fig. 1). Quantification of color classification accuracy, as separately determined using fixed cells singly labeled by CF568 (Fig. 3f) and AF647 (Fig. 3g), indicated that at ~5000 photons, excellent accuracies of 98.2% were achieved for both dyes at the confidence threshold of 0.8. At ~3000 photons, the accuracy for CF568 did not vary noticeably (Fig. 3f), whereas the accuracy for AF647 dropped to ~90.4% (Fig. 3g). Lowering the confidence thresholds led to accuracy drops by a few percentage points (Fig. 3fg). Previous work^23^ has shown that for dyes in these two color channels, through traditional sequential imaging using different optical filter sets, a ~8% crosstalk occurred from the 561-nm excited dye into the 647-nm excited dye, whereas crosstalk in the opposite direction was ~1%. Our accuracies thus appear to outperform at ~5000 photons, a value often obtained in STORM experiments^9,23^. Moreover, in our case, all data were collected within the same optical path in a single STORM session, so we avoided the major difficulties in aligning images from different filter sets.

Based on our successful color classification, two axial-localization ANNs, each trained for AF647 and CF568, were next used to separately decode the axial positions of the molecules in the two color channels, the results of which were recombined into one image for presentation (Fig. 3e and Supplementary Fig. 1). This showed the expected result that the cell edges, thinner in height, were dominated by small z values, whereas for regions far away for the cell edges, the cells became thicker and had increased z values. White arrows in Fig. 3ae further point to regions of microtubules, labeled by the same CF568 dye, where similarly increased PSF widths were noted, but the ANN correctly identified one being below the focal plane whereas the other above. We further note that as the two color channels are successfully separated, they may also be separately fed into other recent methods that extract axial positions from unmodified PSFs^17,18^.

As a final note, our end-to-end framework is further characterized by exceptional speed. Once trained, evaluation speeds were >3.3×10^5^ molecules/s (GPU acceleration) for both the color-separating and the axial-localization ANNs. These speeds surpass traditional “fitting” approaches^18^, and so could be especially beneficial for speed-demanding applications including online analysis.

Our finding highlights the rich, multidimensional information concealed in the details of the diffraction-limited image of a fluorophore, which was uniquely unleashed in this work through state-of-the-art machine learning algorithms. Not having to modify the PSF shape or divide single-molecule fluorescence between different optical paths, or to image sequentially, not only simplify experimental implementation, but more importantly, preclude the deterioration in SMLM performance due to enlarged PSFs and/or split channels, as well as the need to align localizations from different channels. Meanwhile, we note that our approach is not limited to a particular type of PSF or machine-learning algorithm. The co-evolution of our data-driven end-to-end framework with the ongoing efforts on PSF engineering^8^ should thus lead to new improvements, and conceivably, fundamentally new types of imaging modalities, for multidimensional SMLM.

---

**Supplementary Fig. 1.**
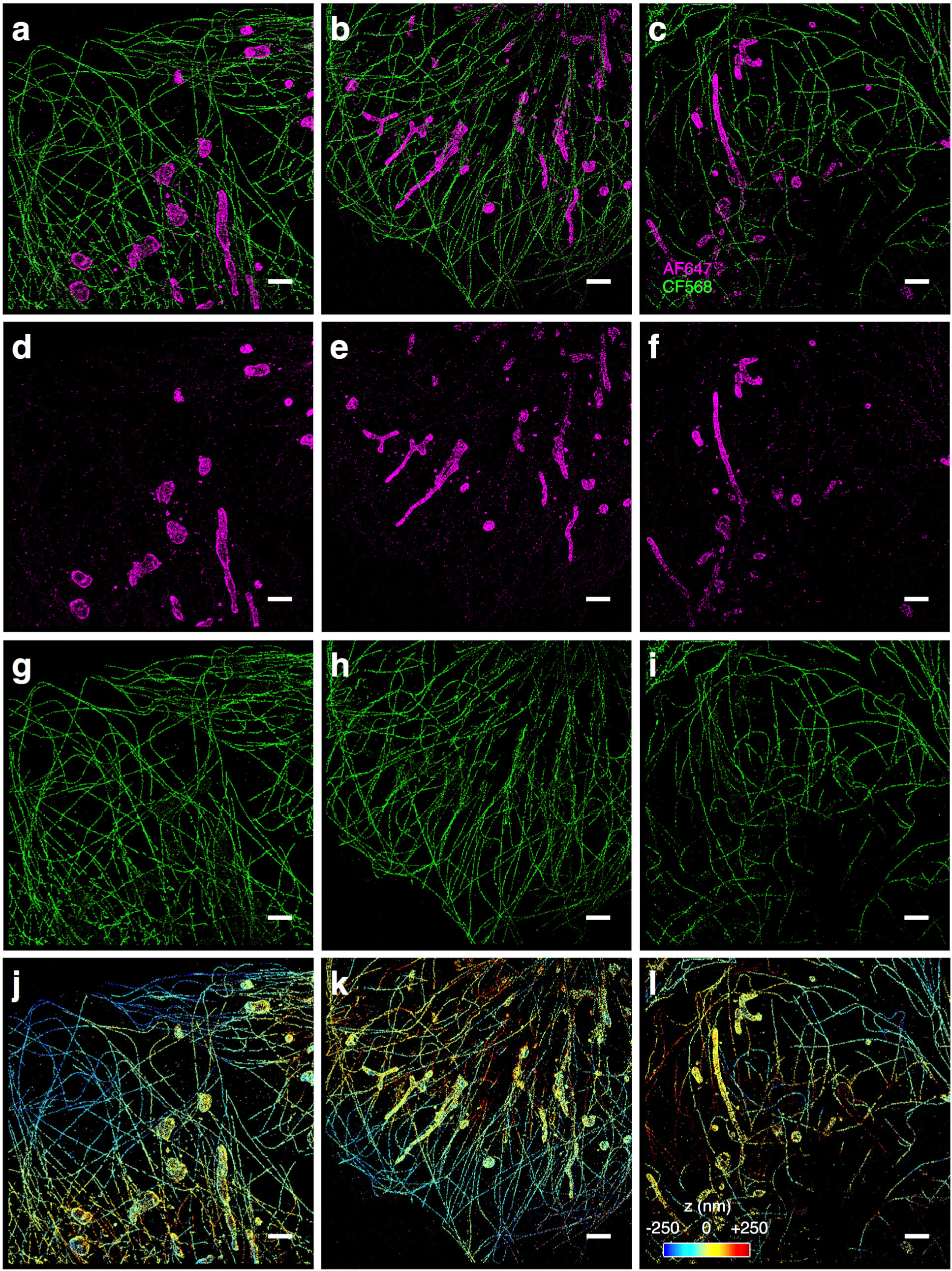
Additional examples of ANN-resolved multicolor 3D SMLM of fixed cells based on unmodified PSFs. Scale bars, 2 μm.

## Methods

### Optical setup

STORM and bead experiments were performed on a Nikon Ti-E inverted fluorescence microscope using an oil-immersion objective lens (Nikon CFI Plan Apochromat λ 100×, NA 1.45) and the native 1.5x magnification on the microscope, without any modifications to the imaging path. Lasers emitting at 644, 561, and 488 nm were introduced to the back focal plane of the objective lens via a multi-line dichroic mirror (ZT405/488/561/640rpc-uf2, Chroma). A translation stage shifted the laser beams toward the edge of the objective lens so that they entered slightly below the critical angle, illuminating <1 µm into the sample. Emission was filtered by a multi-notch filter (ZET405/488/561/640m, Chroma) and recorded by an EM-CCD camera (iXon Ultra 897, Andor). Effective magnification and pixel size were ~150x and ~107 nm, respectively.

### Bead samples

40 nm dia. fluorescent beads from Invitrogen (F10720; “yellow” and “orange” FluoSpheres with emission peaks at 515 and 560 nm, respectively) were diluted in Dulbecco’s phosphate buffered saline (DPBS), mixed, and sealed between a glass slide and a pre-cleaned #1.5 thickness coverslip, and imaged with the above optical setup. The 488-nm and 561-nm lasers were used to excite the two types of beads to similar levels of brightness. To record images at different axial positions, the objective lens was scanned by the built-in motor over a range of −200 to +200 nm of the focal plane.

### Cell samples

COS-7 cells were plated on #1.5 coverslips to reach a confluency of ~50% in ~1.5 days, and fixed with 0.1% glutaraldehyde and 3% paraformaldehyde in DPBS at room temperature. The sample was quenched with 0.1% sodium borohydride in DPBS and rinsed with DBPS three times. Primary and secondary antibodies were diluted in a blocking buffer (3% BSA + 0.1% Triton X-100 in DPBS) and labeled as described previously^9^. Primary antibodies were mouse anti-tubulin (Abcam ab7291) for microtubules and rabbit anti-Tom20 (Santa Cruz sc-11415) for mitochondria. Secondary antibodies were AF647-labeled goat anti-mouse IgG1 (Invitrogen A21240), AF647-labeled goat anti-rabbit IgG (Invitrogen A21245), and donkey anti-mouse IgG (Jackson ImmunoResearch) conjugated with a CF568 succinimidyl ester (Biotium 92131). Samples for training the color-separating ANN were singly labeled for microtubules with CF568 or AF647, whereas for the two-color unknown samples, microtubules and mitochondria were respectively labeled with CF568 and AF647. The sample was mounted in a STORM buffer [10% (w/v) glucose, 120 mM cysteamine, 0.8 mg/mL glucose oxidase, and 40 µg/mL catalase in Tris-HCl (pH 7.5)] and imaged using the optical setup described above. For consistent experimental conditions, all cell samples were imaged at comparable depths with the focal plane being ~300 nm away from the coverslip surface. The sample was illuminated with the 561 and 644 nm lasers at ~2 kW/cm^2^, which led to the photoswitching of CF568 and AF647 single molecules. Fluorescence was recorded by the EM-CCD for a frame size of 256×256 pixels at 55 frames per second. Each movie was typically recorded for 20,000 frames.

### Antibody samples

Training data for the axial localization of AF647 and CF568 single molecules were acquired using the above AF647- and CF568-labeled secondary antibodies. The two antibodies were separately diluted in DPBS to ~2 µg/mL. Pre-cleaned #1.5 coverslips were separately incubated in either solution for ~5 min, briefly air-dried, rinsed with distilled water, and mounted and imaged as described above. To record single-molecule images at different axial positions, the objective lens was scanned by the built-in motor in the range of −700 to +700 nm of the focal plane.

### Preprocessing of single-molecule images for ANNs

Single-molecule fluorescence in raw STORM and bead data was first identified and localized in 2D using established methods^1^. Single-molecule PSF images were cropped as 13×13 pixels surrounding the 2D localizations; at this time, images containing multiple emitting molecules were rejected. The cropped PSF images were zero-centered, and their L2/Euclidean norm was normalized before being used as inputs for ANNs.

### Design and implementation of neural networks

An ANN architecture comprising multiple hidden layers was implemented using the Tensorflow framework on a computer with 32GB RAM, Intel i7-7800X CPU, and Nvidia GTX-1080Ti GPU. The same architecture was used for both the color-separating and axial-localization ANNs (4 total layers of 4096-4096-2048-1024 neurons, respectively). Each hidden layer was fully connected, and rectified linear units were used as their activation function. For color discrimination, Softmax function and cross-entropy were used for loss calculation, the weights in the network were not directly included for regularization, and a dropout layer was inserted before the final layer to prevent over-fitting. For axial localization, the output of the final layer was set to be a scalar value, and L2 norm was used to calculate the learning loss for each batch. In this case, L2 norm of the weights in each layer was added to the loss function for regularization, and dropout was not used. Network hyper-parameters such as the number of neurons in each layer (given above), the dropout ratio (0.5 for the color-separating ANN), and the regularization factor (0.01 for the axial-localization ANNs) were adjusted for optimized performance.

### Neural network training

Since the network is subject to handling input images with various noise levels, it was essential to maintain a consistent noise level within the training dataset regardless of the classification class or axial location. Therefore, experimental PSF images with comparable photon counts of 4500-5500 were used throughout the training process, and molecules with photon counts higher than this range were used for the validation during the training process to check the generalization of the trained network. The weights initialized by the Xavier method^24^ are trained using the Adam optimization algorithm. An initial learning rate of 10^−4^ and 10^−3^, and a batch size of 64 and 32 were used for the color-separating and the axial-localization ANNs, respectively. The learning rate was set to decrease by ~5x after every 1,000 iterations. The sizes of the training sets were ~10,000 and ~6,000 per fluorophore type for the color-separating and the axial-localization ANNs, respectively. The networks converged within ~10 epochs (training time: 231.9 s for the color-separating ANN, and 488.5 s for the axial-localization ANNs).

### Neural network inference

At the inference stage, input single-molecule PSF images were first plugged into the color-separating ANN. This ANN provides a conditional probability distribution corresponding to the input image. Specifically, when the size of each input image is *N* by *N* pixels, and there are *M* different molecule color classes, the final output from the Softmax layer for the *i*th input image is

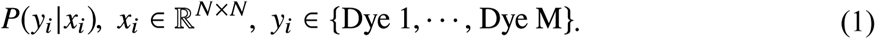

From this distribution, the ANN makes the decision in a maximum-likelihood manner: the molecule color class with the highest probability is chosen. This, in turn, implies that we can use this distribution to quantify the classification confidence. For example, in a simple binary classification problem, the confidence for the color assignment of the *i*th input image can be evaluated as

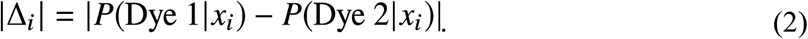

By setting a finite confidence threshold *δ* to reject molecules with low classification confidences (|*Δ*^*i*^|<*δ*), improved classification accuracy may be obtained (Figs. 2ef and 3fg). This parameter may thus be adjusted by the user to balance the classification accuracy and rejection rate. Once the color of the molecule is determined, the single-molecule image is plugged into the axial-localization ANN trained for that particular color to evaluate the axial position. With GPU acceleration, both inference steps (passing the forward path of the neural networks) were extremely fast: only ~300 ms was used to infer 100,000 molecules for both the color-separating and the axial-localization ANNs.

## Acknowledgments

This work was in part supported by the Beckman Young Investigator Program and the Packard Fellowships for Science and Engineering. K.X. is a Chan Zuckerberg Biohub investigator. T.K. acknowledges support from Kwanjeong Educational Foundation. S.M. acknowledges support from Samsung Scholarship.

